# A practical and robust method to evaluate metabolic fluxes in pancreatic islets

**DOI:** 10.1101/2023.07.25.550538

**Authors:** Debora Santos Rocha, Antonio Carlos Manucci, Alexandre Bruni-Cardoso, Alicia J. Kowaltowski, Eloisa A. Vilas-Boas

## Abstract

**Aims/hypothesis:** Efficient mitochondrial oxidative phosphorylation is essential for pancreatic beta cell responses to nutrient levels. Consequently, the evaluation of mitochondrial oxygen consumption and ATP production is important to investigate essential aspects of pancreatic islet pathophysiology. Currently, most studies use cell lines instead of primary islets due to difficulties in measuring primary islet respiration, which requires specific equipment and consumables that are expensive, complicated to use, and poorly reproducible. The aim of this study is to establish a robust and practical method to assess primary islet metabolic fluxes using Extracellular Flux Technology and standard commercial consumables.

**Methods:** Pancreatic islets were isolated from 8 to 12-week-old mice and rats, and submitted to a dispersion protocol using trypsin. Dispersed islets were adhered overnight to pre-coated standard Seahorse microplates, and oxygen consumption rates were evaluated using a Seahorse Extracellular Flux Analyzer. We also validated the functionality of dispersed islets by analyzing glucose-stimulated insulin secretion (GSIS) and calcium (Ca^2+^) influx in response to different modulators by fluorescence microscopy.

**Results:** We provide a detailed protocol with all steps necessary to optimize islet isolation and dispersion, in order to achieve a high yield of functional islets and perform metabolic flux analysis. With this method, which requires only a few islets per replicate, both rat and mouse islets present robust basal respiration and proper response to mitochondrial modulators (oligomycin, CCCP, antimycin and rotenone) and glucose addition. Both oligomycin and CCCP concentrations were titrated. Our method was also validated by other functional assays, which show these cells present conserved Ca^2+^ influx and insulin secretion in response to glucose.

**Conclusions/interpretation:** We established a practical and robust method to assess *ex vivo* islet metabolic fluxes and oxidative phosphorylation. Our findings cover an important gap in primary islet physiology studies, providing a valuable tool we hope is useful to uncover basic beta cell metabolic mechanisms, as well as for translational investigations, such as pharmacological candidate discovery and islet transplantation protocols.

**Research in context:** *What is already known about this subject?:*

- Pancreatic beta cells efficiently couple oxidative phosphorylation and ATP production with insulin secretion; mitochondrial ATP production is crucial for proper insulin secretion.
- Most studies of beta cell respiration use cell lines instead of primary islets, which are a much more robust model to evaluate beta cell function.
- The few works with primary islet respiration use specific equipment and consumables that are expensive, complicated, and poorly reproducible.

*What is the key question?:*

- Is it possible to develop a practical method to evaluate metabolic fluxes and ATP production in isolated islets, using the standard Seahorse Extracellular Flux Technology?

*What are the new findings?:*

- We optimized rodent islet isolation and functional analysis protocols using standard extracellular flux analysis equipment and consumables.
- Our method allows for increased islet yield and robust islet respiration measurements.

*How might this impact on clinical practice in the foreseeable future?:*

- Quantitative measurements of metabolic fluxes and oxidative phosphorylation are the cornerstone of new discoveries in beta cells, and can contribute toward the establishment of new cellular protocols, such as for cell transplantation, as well as the development of new pharmacological agents targeted to these cells.

## 1. Pancreatic beta cells and mitochondrial respiration

The endocrine pancreas is composed by pancreatic islets, highly vascularized micro-organs formed by aggregates of distinct cell types, each responsible for the production and release of a different hormone: glucagon-producing alpha cells, insulin-producing beta cells, and somatostatin-producing delta cells, among others. The average diameter of a whole human islet is similar to the average diameter of a rat islet: 108 ± 6 and 115 ± 5, respectively [1]. However, islet sizes vary largely within samples from the same animal, ranging from a few endocrine cells to a large islet composed of several thousand cells [2]. In addition, the percentage of each cell type within the islet and the spatial arrangement of the different cell types within the islet also vary among species; beta cells are normally the predominant cell type [3, 4].

Pancreatic beta cells are responsible for insulin synthesis and release in response to increased postprandial plasma nutrients, especially glucose, which is the main insulin secretagogue. Glucose-stimulated insulin secretion (GSIS) control [5] begins with the entry of glucose into beta cells through specific glucose transporters (GLUTs). Glucose, if present at levels sufficient to achieve affinity, is then rapidly phosphorylated by glucokinase, generating glucose-6-phosphate, which proceeds through the glycolytic pathway, generating pyruvate. Pyruvate then enters mitochondria, where it is converted into acetyl-CoA and completely oxidized by the tricarboxylic acid (TCA) cycle. The TCA cycle and oxidative phosphorylation efficiently generate ATP in beta cell mitochondria, and the increased cellular ATP/ADP ratio leads to closure of ATP-sensitive potassium channels (K_ATP_) at the plasma membrane, with subsequent plasma membrane depolarization. This promotes opening of voltage-sensitive Ca^2+^ channels in the plasma membrane, inducing Ca^2+^ influx, activating exocytotic machinery, and culminating in insulin release.

Since pancreatic beta cells efficiently couple glucose metabolism to insulin release through mitochondrial electron transport and ATP production [6, 7], studying oxygen consumption rates (OCR) in *ex vivo* islets is of great interest in the areas of beta cell physiology and metabolic regulation. Indeed, mitochondrial respiration and insulin secretion may be altered under pathological conditions such as type 1 and 2 diabetes [8, 9], making the analysis of mitochondrial respiration also important for islet functional assessment in the context of drug discovery and islet preservation protocols.

Different methods have been developed to measure OCRs in different tissues and cell types. Measurements can be done in suspended isolated mitochondria, as well as suspended intact or permeabilized cells using an Oroboros high-resolution oxygraph. In plated intact cells, OCRs can be measured with high sensitivity using a Seahorse Extracellular Flux system; however, most current studies of beta cell respiration use beta cell lines, instead of primary islets. The few studies with primary islet respiration published so far [10–16] use specific equipment and consumables that are expensive, complicated to use, and that yield results that may be poorly reproducible. While cell lines are also an important tool, primary islets are a much more robust model to evaluate beta cell function.

Here, we briefly discuss methods available to measure isolated islet oxygen consumption and their limitations. In order to make oxygen consumption more readily measurable in primary islets, we describe a new and practical protocol to assess isolated pancreatic islets using Extracellular Flux Technology and standard microplates. We optimized the technique for both mice and rats, and found that it results in increased islet yield and robust reproducibility.

## 2. Existing methods to measure oxygen consumption in islets

Islet respiration has been measured using Oroboros high-resolution oxygraphy, where islets are suspended in a medium/buffer and their oxygen consumption rates are followed over time. This is an indisputably robust method to measure OCRs, suitable for many cell types and tissues. However, there are two main disadvantages in the case of islets. Measurements are made in chambers that support 0.5 mL or 2 mL of media. These large volumes require a high number of cells, which is an important limitation for islets.

Islet isolation is a multi-step protocol. The yield depends largely on the animal species used and training of the investigator, but since islets are around 2% of all cells in the pancreas, the yield is normally low. In our hands, basal respiration levels were very low using up to 80 rat islets in a 2 mL chamber. Indeed, the required number of islets seems to be much higher, around 250 mouse islets per mL [10], which results in large use of animals, as one mouse yields approximately 200 to 350 islets, while one rat yields approximately 400 to 600 islets. In addition, constant stirring is necessary during oxygen electrode measurements, to ensure proper oxygen diffusion. This can damage the islets, leading to dispersion of the cells and decreased viability [11]. Specific devices have been developed to overcome this, such as a 3D-printed chamber to stabilize the islets during Oroboros measurements [11]. As an alternative to Oroboros, other strategies have also been developed, such as an islet-on-a-chip microfluidic device that allows OCR measurements at the same time as calcium (Ca^2+^) imaging of individual islets [12]. However, these devices are not readily available commercially.

The Seahorse Extracellular Flux system, widely available in most major research centers, was designed specifically for adherent cells, and has enhanced sensitivity while also avoiding stirring damage. However, the spheroid arrangement of islets is not adequate for use in its standard plate-based systems, in which the sensors are located close to the adherent plate surface during measurements, to enhance sensitivity.

There are specific, commercially available, Seahorse 96-well microplates for spheroids, with round bottoms which could favor placing islets in the center. However, working with 3D structures can be tricky, and several adjustments and precautions must be taken into account depending on the cell type used [17]: some researchers place the islets under a microscope to orient pipetting [13], centrifuge islets after pipetting [13, 14], and in all cases pre-coating of the plates is required for the islets to properly adhere to the bottom and avoid movement during measurement [13–15]. Even with these precautions, islets may still detach during mixing and measuring steps. In addition, a special thermal tray compatible with the spheroid microplate is required for the measurements, so the equipment must be reserved for spheroid plates, and cannot interchangeably use other plates adequate for most adherent cell types.

In order to circumvent these difficulties, and try to keep islets trapped at the bottom of the well while minimizing variabilities, special Seahorse 24-well microplates were specifically designed for use with islets [16]. Islets are placed in a depression at the bottom of the well with a special tool (Islet Capture Screen Insert Tool), and each well has a grid of polycarbonate to lock in the islets and keep them from detaching during mixing and measuring steps. Of note, the spheroid microplates can only be used in 96-well Seahorse systems, while islet capture microplates can only be used in 24-well Seahorse systems, limiting the choice of use for researchers who have access to only one of these specialized analyzers.

Overall, it seems clear to us that several limitations that hinder robust and reproducible OCR assessment as a measure of mitochondrial activity in primary islets persist today, and are resulting in limited measurements of this important functional parameter. These limitations are mostly due to the high number of islets required for the measurements, loss of viability during the measurements, special devices and consumables that are tricky to use and expensive, in addition to poor reproducibility between replicates. In the next sessions, we present a detailed protocol of a practical method that allows robust and reproducible OCR measurements using a relatively low number of islets and standard materials.

## 3. Optimization of rodent islet isolation

We isolated pancreatic islets from C57BL/6NTac male mice and Sprague Dawley female rats. Animals were bred and housed in the animal facility of the Faculty of Pharmaceutical Sciences and Chemistry Institute of the University of São Paulo, devoid of murine specific pathogens. Animals were maintained in collective cages (max 5/cage) at controlled temperatures (23°C) in a 12-12 h light/dark cycle with free access to food/water. We used 8 to 12-week-old animals, randomly selected for the experiments. Rats weighed 280 to 350 g and mice 25 to 30 g. All procedures were conducted in accordance with the Ethical Principles of Animal Manipulation of the local animal ethics committee, under protocols CEUA-IQ/USP 196/2021 and 244/2022.

### 3.1. Pancreatic islet isolation

Pancreatic islet isolation is of great importance to study endocrine physiology and pathology, especially in type 1 and 2 diabetes. Lacy et al. in 1967 [18] originally developed a protocol using a collagenase solution in Hanks buffer, and highlighted a first crucial point for this method: properly clamping the Ampulla of Vater region, the meeting point at the duodenum of the bile and pancreatic ducts. This prevents leakage of the collagenase solution into the intestine, avoiding inadequate inflation of the pancreas, and allowing visualization for total excision of the tissue and yield optimization [18]. Since this method was first proposed, the protocol has been optimized, leading to even greater islet yields per animal. Corbin et al. [19] highlighted the importance of using adequate collagenase for exocrine tissue digestion. Type V, XI, and P collagenases are the most suitable, significantly impacting on yield and viability.

In the protocol proposed in this paper, we use type V collagenase diluted in Hanks’ buffer (0.7 mg/mL), which was filtered using a 0.2 µm filter to avoid the infusion of particles, and consequent physical damage to the tissue, in addition to avoiding contamination in culture. Collagenase solutions were kept on ice until infusion through the common bile duct. Using this protocol, after 25 minutes digestion of the exocrine tissue at 37°C in a water bath, we recovered at least 250 islets per mouse and 400 per rat. Immediately after digestion, it is important to keep the solution on ice to decrease proteolytic activity and prevent digestion and death of the endocrine tissue.

Centrifugation, discarding the supernatant, and washing with a fresh buffer are additional crucial steps in the protocol. Still in 1967, Lacy et al. [18] compared the effects of centrifugation after digestion to simple sedimentation by gravity, leading to an average yield gain from 100 to 300 islets per mouse, close to the number obtained with our protocol after digestion. However, they also pointed out that sucrose gradient centrifugation, not used here, may be harmful to the functionality of the islets.

Salvalagio et al. [20] compared the use of a Ficoll gradient and cell strainer filtration in regard to the yield and functionality of the islets. The use of a Ficoll gradient was less effective in comparison to filtration when evaluating the number of islets, GSIS, and purity. Also, islets isolated by filtration were able to restore normoglycemia when transplanted into diabetic mice one day after the procedure, while islets isolated by Ficoll need 6-7 days to promote the same result. Regardless of the method used, the authors point out that manual collection significantly increases the purity of the islets. In the procedure described in our paper, the collection of the islets was done manually, a method that is still advantageous in terms of the functionality of the islets compared to gradient centrifugation.

It is noteworthy that all steps involved in islet isolation are relatively simple, but require practice by the researchers involved. Like any surgical procedure, prior practice implies better effectiveness and reduces the time spent on the procedure [21]. Thus, a pilot experiment is mandatory to ensure a reasonable average number of islets and to avoid discarding animals due to lack of practice. Importantly, the time between the death of the animal and the separation of the endocrine and exocrine tissues has great impact on maintaining the viability and functionality of the islets. Thus, when using more than one animal per day, kill one at a time and isolate the tissue as freshly as possible. Finally, the age of the animals also impacts on yield, with higher yield values being observed in older animals compared to juvenile animals [19].

At the end of the isolation protocol, the islets can be easily visualized on a dark background plate (Fig. 1B), without staining.

**Figure 1.**
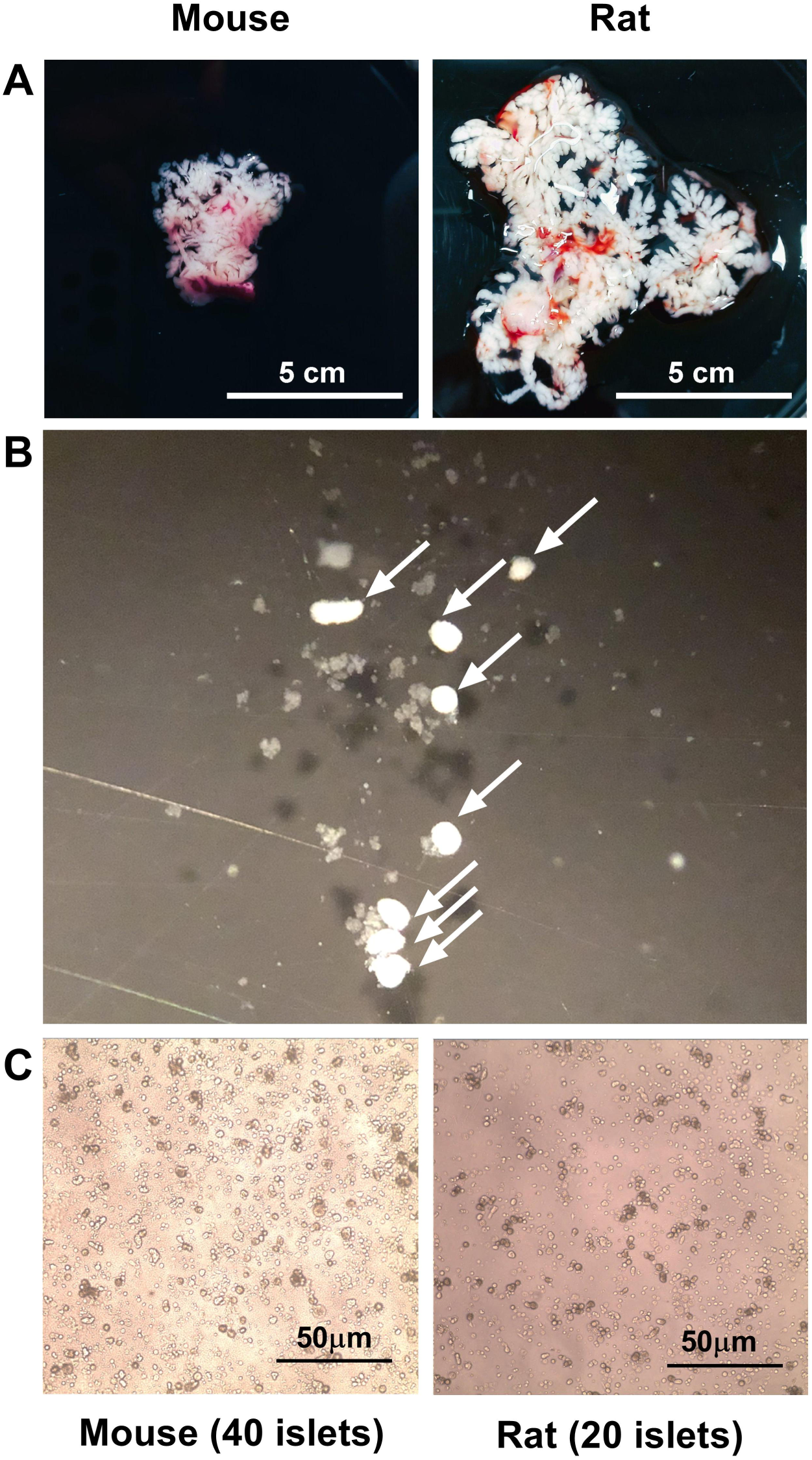
Pancreas isolation, islet dispersion, and culture. **(A)** After pancreas dissection and full inflation, whole mouse and rat pancreases can be visualized macroscopically. **(B)** After exocrine digestion with trypsin, isolated islets can be visualized using a stereomicroscope. **(C)** Dispersed and adhered islet cells in culture can be visualized using a microscope after hand-picking, dispersion with trypsin, and overnight adhesion on a pre-coated microplate. Representative figures are shown.

### 3.2. Pancreatic islet isolation - Optimized protocol

1. Anesthetize the animals as recommended by the local ethical committee. We used 70 mg/kg ketamine, 10 mg/kg xylazine, and 2 mg/kg acepromazine, followed by cervical dislocation (mice) or cardiac puncture (rats). **Caution:** We obtained an islet yield ranging from 100 to 250 per mouse and 200 to 350 per rat. Importantly, islet yield increases in freshly isolated pancreases. Thus, when using more than one animal per day, it is recommended to anesthetize one at a time. In addition, islet collection (and separation from the exocrine tissue) must be performed readily. As islet collection is a time-consuming technique, we used a maximum of two rats / three mice per day per investigator.
2. With the animal in *dorsal decubitus*, clean the abdominal area with 70% ethanol, and access the abdominal cavity. **Caution**: Islets tend to stick to furs, so use 70% ethanol to wet the animal and keep surgical material clean. **Note:** Due to the small size of the mice and their respective ducts, we recommend using a stereomicroscope during steps 3 and 4.
3. Place the animal’s head close to the surgeon and still under *dorsal decubitus*. Access the abdominal cavity and locate the common bile duct. Clamp the end of the Ampulla of Vater using a Spencer Wells forceps to prevent collagenase from going into the intestine. **Note:** Clamping the Ampulla of Vater with a surgical line is also feasible.
4. Make a small cut in the duct, not transposing it, in order to insert a needle (30G) and infuse it with 5 mL (mice) or 15 mL (rat) of cold collagenase type V in Hanks’ Balanced Salt Solution (HBSS, table 1) (0.7 mg/mL). **Caution**: The collagenase solution must be sterile and at 4-8°C **Caution:** To avoid contamination in the culture and physical damage to the tissue, the collagenase solution must be filtered (0.2 µm filter) before inflating the pancreas. **Caution**: Avoid bubbles and do not inject air into the tissue.
5. After full inflation, remove the whole pancreas carefully, to prevent it from breaking or allowing intestinal content to pour into the tissue. **Note:** Fig. 1A shows a whole mouse and rat pancreas after full inflation. **Caution**: Avoid spending more than 15 min per animal on the pancreas inflation procedure.
6. Mince the tissue into small fragments using sharp scissors. **Caution**: Maintain the tissue-collagenase mixture at 4-8°C if using more than one animal at a time.
7. Incubate the pancreases for 25 min at 37°C (water bath) under gentle shaking for digestion of the exocrine tissue.
8. After incubation, the pancreas should be shaken manually, not vortexed, until the tissue is clearly dissociated in a homogenous solution with no more than a few small pieces of tissue.
9. Place the tube containing the homogenous solution immediately on ice to stop the digestion.
10. Complete to a final volume of 20 mL with HBSS (4-8°C) (Table 2).
11. Centrifuge at 1,000 rpm (130 g) for 3 min at 4°C.
12. Carefully discard the supernatant, keeping a small part of the liquid phase to avoid tissue loss, and re-suspend the pellet with 20 mL of ice-cold HBSS buffer.
13. Steps 11 and 12 should be done thrice.
14. Put the solution in a dark background Petri dish, and collect islets manually to another plate with cold HBSS using a micropipette under a stereomicroscope, keeping everything, other than the stereomicroscope plate, on ice. **Note:** Fig. 1B shows the isolated islets in Hanks’ buffer after handpicking, with a few digested exocrine tissue pieces. The arrows indicate the isolated islets, which are round, white structures surrounded by digested exocrine tissue. **Note:** Staining the islets with special dyes is also possible to facilitate visualization of the islets, but it is not necessary. **Note**: Ficoll, Percoll and sucrose gradients can also be used for purification, but islet viability and functionality may be lower when compared to handpicking.

**Table 1.** - Comparison of different Seahorse-based methods.

|  | **Islet microplate** | **Spheroid microplate** | **Dispersed cell measurements** |
| --- | --- | --- | --- |
| **Compatible system** | Seahorse XF24 | Seahorse XF96 | Seahorse XF24 and XF96 |
| **Microplate** | Specific islet microplate | Specific spheroid microplate | Standard microplate |
| **Additional consumables required** | Islet capture screen insert tool | Specific thermal tray for spheroid | No |
| **Islets per condition** | 50 - 80 (rat or mouse) | 10 - 20 (mouse) | 20 (rat)<br>40 (mouse) |
| **Precoating required** | No | Yes | Yes |
| **Whole islet / dispersed cells** | Whole islets | Whole islets | Dispersed cells |
| **Centrifugation required** | No | Yes | No |
| **Allows microscopy** | No | No | Yes |

**Table 2.** - Composition of Hanks’ Balanced Salt Solution (HBSS).

| **PART 1** |  |  |
| --- | --- | --- |
|  | **g/L** | **mM** |
| **NaCl** | 8.18 | 140 |
| **KCl** | 0.37 | 5 |
| **MgSO<sub>4</sub>** | 0.048 | 0.4 |
| **Na<sub>2</sub>HPO<sub>4</sub>** | 0.017 | 0.1 |
| **KH<sub>2</sub>PO<sub>4</sub>** | 0.054 | 0.4 |
| **PART 2** |  |  |
|  | **g/L** | **mM** |
| **CaCl<sub>2</sub>** | 0.0838 | 0.75 |
| **NaHCO<sub>3</sub>** | 0.336 | 4 |
| **D-glucose** | 1.8 | 10 |
| **BSA** | 2 | 0.2 % |
| **P/S** | 10 mL | — |

**Table 3.**
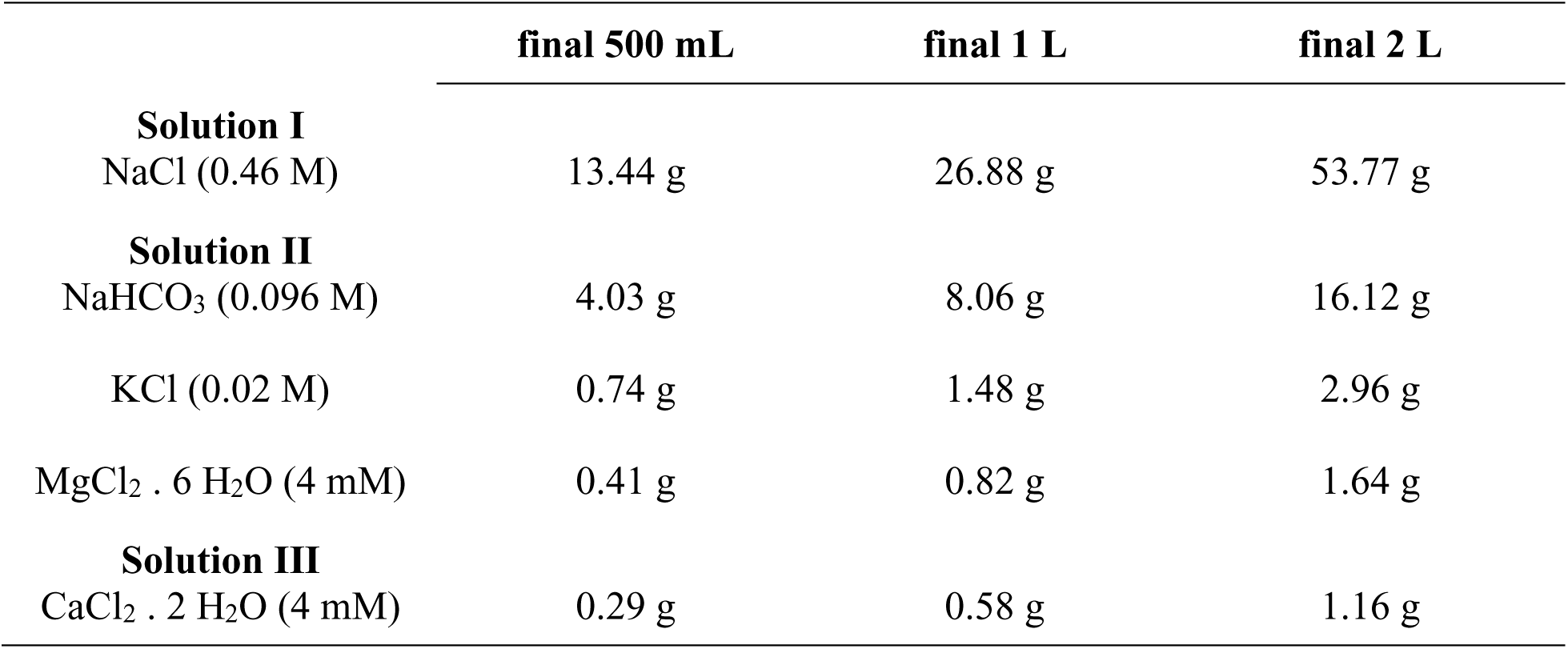
- Composition of Krebs Henseleit.

### 3.2. Pancreatic islet dispersion and culture

An important limitation of using intact islets is that large islets often display insufficient oxygen diffusion toward the islet core, leading to central necrosis due to hypoxic conditions [22, 23]. This situation may be aggravated when keeping islets for days in culture. In this paper, we chose to use dispersed islets adhered to standard flat-bottom Seahorse XF24 microplates. The use of dispersed cells to assess islet function is not new, and different protocols using trypsin, accutase or mechanical dispersion have been used by several authors [13, 24, 25]. It is well known that the three-dimensional structure of the pancreatic islet and the interaction between different endocrine cell types (both *in vivo* and *ex vivo*) is important for its function. Thus, paracrine interactions have an impact on environmental responses [26, 27]. Therefore, it is important to point out that, although most dispersion protocols are relatively fast and simple, validation using functional protocols may be necessary to guarantee adequate functionality and viability if interactions between cells are expected to participate in the biological phenomena studied. This was a concern we had throughout our study, as we describe in further sessions (item 3.4).

#### 3.2.1. Pancreatic islet dispersion and culture - Protocol

1. After handpicking the islets, collect 20 (rats) or 40 (mice) islets in conic microtubes containing 300 µL cold trypsin-EDTA solution (0.25%). **Note:** There is a natural tendency to choose the best islets first, so to minimize variations between replicates of the same day, we collected the first same-sized 20 islets and distributed them in all tubes first, and then collected the final same-sized 20 islets.
2. Incubate the solution (islets in trypsin) at 37°C for 2 min with gentle shaking.
3. Place the tubes immediately on ice and inactivate the trypsin with 600 µL complete RPMI medium (with 10% fetal bovine serum, FBS).
4. Centrifuge at 1,000 rpm (130 g) for 1 min to pellet the cells.
5. Discard the supernatant carefully with a micropipette and re-suspend the dispersed islets in 150 µL complete RPMI medium. **Note:** If necessary, a spotlight can be used to visualize the pellet better and avoid discarding the islets along with the supernatant.
6. Homogenize the pellet by pipetting up and down until complete visible dispersion.
7. Transfer the whole volume to cell culture plates previously coated with poly-L-lysine (as described at 3.3).
8. Let the cells adhere overnight at 37°C in a humidified atmosphere of 5% CO_2_ / 95% air. **Note:** Fig. 1C shows dispersed cells from mouse (left) and rat (right) pancreases.

### 3.3. Poly-L-lysine coating

Proper cell adhesion to the wells is of extreme importance, to avoid loss of cell adhesion during mixing and measuring steps. Here, all cell culture microplates were coated with poly-L-lysine (#P8920, Sigma) prior to use. We found that letting the cells adhere overnight, rather than for a few hours, ensures proper adhesion and reduces variability between replicates.

1. Using sterile material and reagents, add 50 µL/well of prediluted poly-L-lysine (final 0.001%) solution to the Seahorse XF24 microplate and incubate for at least 1 h at 37°C.
2. Rinse wells twice with 200 µL/well of sterile milliQ water.
3. Discard the water and let the wells dry at room temperature inside a laminar flow cabinet.
4. After completely dry, seal the plates with parafilm and stock at 2-8°C for up to 10 days.

### 3.4. Validation of the protocol by functional assays

As previously mentioned, since the main aim of the proposed protocol is to evaluate pancreatic islet metabolic fluxes in physiology and pathology, the validation of the functionality of these islets after the dispersion protocol and adherence to the culture microplate is mandatory.

In 1984, Weir G.C. et al. [24] used Sprague-Dawley rat islets dispersed with 0.1% trypsin for 15 min. The authors compared their functionality immediately after dispersion and after 2-3 days of adhesion in culture. Immediately after dispersion, the cells did not respond to glucose in a GSIS assay. However, when the dispersed cells were allowed to adhere to the culture microplate, the authors observed proper insulin secretion in response to glucose. Glucagon and somatostatin release upon physiological stimuli were also observed in these cells. Additionally, they found that proinsulin synthesis was even greater in dispersed cells compared to whole islets. These beneficial effects could be due to cell polarization, as observed by electron microscopy, in which mitochondria were located at the bottom of the well, while the secretory vesicles were polarized towards the surface, with greater contact area with the culture medium.

In 2021, Scarl et al. [25] promoted islet dispersion using trypsin, followed by evaluation of single cell intracellular cytosolic calcium levels using the Fura-2 AM probe. Following glucose exposure, the dispersed cells were able to appropriately respond by increasing intracellular calcium influx. Furthermore, the authors observed that cells responded with heterogeneity: different populations of beta cells responded distinctly to increased glucose, with some more sensitive than others.

These studies support the possibility of evaluating physiological events, such as intracellular calcium signaling and insulin release, in dispersed islets. Accordingly, we evaluated Ca^2+^ signaling and insulin release using our dispersion method, as will be described below.

#### 3.4.1. Calcium (Ca_2+_) influx in response to different modulators in dispersed islets

To evaluate whether the dispersed and adhered cells were able to properly secrete insulin in response to glucose, we checked if all the steps involved in this signaling cascade were operational. We modulated different steps in the insulin secretion pathway using physiological (glucose) and pharmacological (glibenclamide and diazoxide) stimuli. As already known (Fig. 2), glucose is taken up by in beta cells by GLUT (mainly GLUT2) independently of insulin, which leads to increased intracellular ATP levels, increases the probability of closing K_ATP_ channels in the cell membrane, modifying the cell membrane potential, opening Ca^2+^ channels sensitive to the membrane potential, and leading to Ca^2+^ influx, necessary for secretory vesicle trafficking to the cell membrane (Fig. 2). Glibenclamide and diazoxide are known pharmacological modulators of plasma membrane K_ATP_ channels. Glibenclamide has a glucose-like effect on Ca^2+^ influx, acting as a channel blocker, inducing membrane depolarization and opening voltage-sensitive Ca^2+^ channels, thereby inducing Ca^2+^ influx. On the other hand, diazoxide is a K_ATP_ channel opener, decreasing Ca^2+^ influx (Fig. 2).

**Figure 2.**
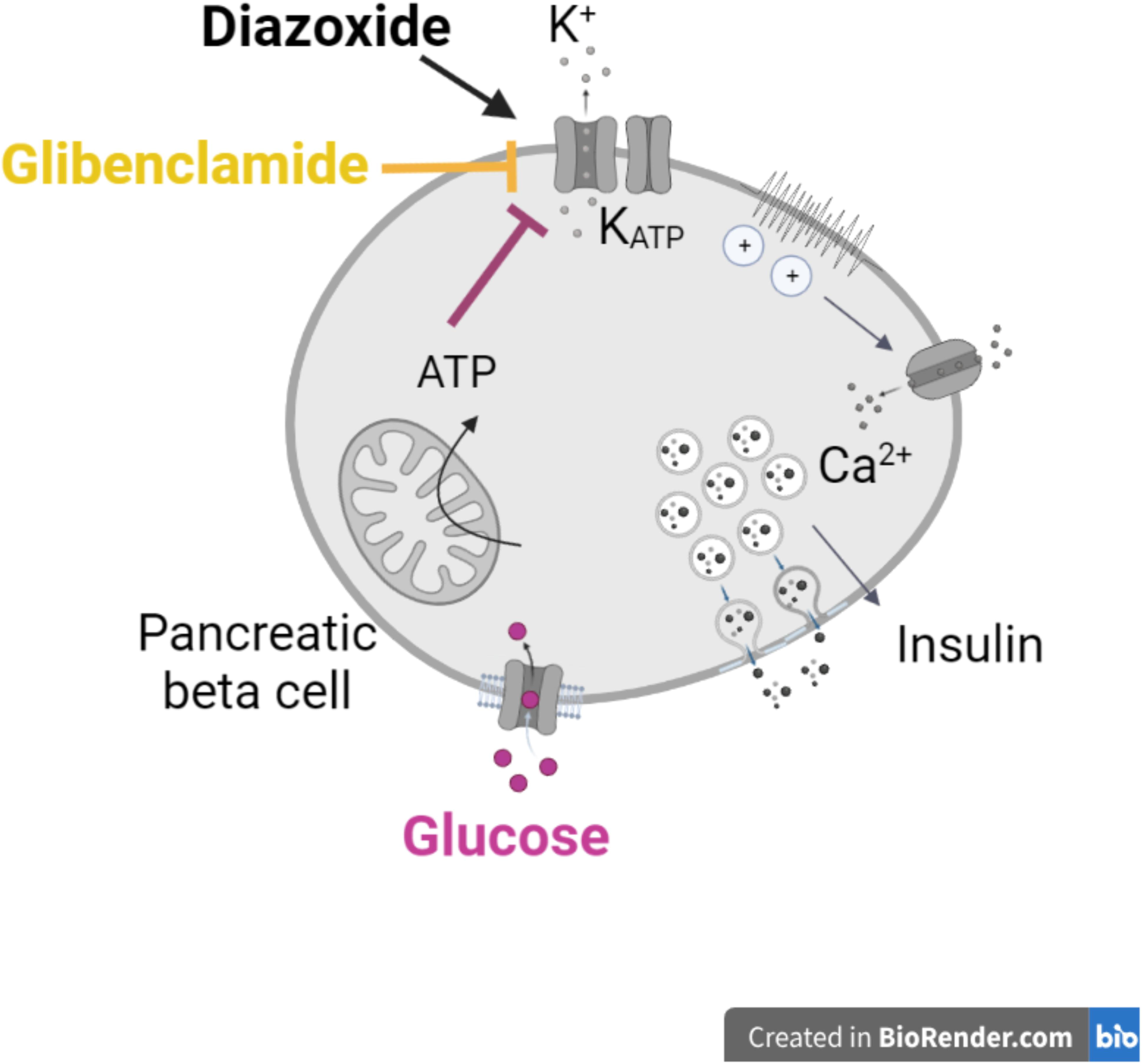
Physiological and pharmacological mechanisms involved in pancreatic beta cell insulin secretion. Glucose **(pink)** is taken up into the beta cell by a specific glucose transporter (GLUT2), independently of insulin. Glucose metabolism leads to increased intracellular ATP levels, and consequently plasma membrane ATP-sensitive potassium channel (K_ATP_) closure and plasma membrane depolarization, voltage-sensitive Ca^2+^ channel opening, Ca^2+^ influx, insulin vesicle trafficking to the cell membrane, and insulin release. Glibenclamide **(yellow)** promotes K_ATP_ closure and diazoxide **(black)**, K_ATP_ opening, leading, respectively, to an increase and a decrease in insulin secretion. Created with BioRender.com.

After overnight incubation on 30 mm cell culture glass dishes previously coated with poly-L-Lysine, dispersed islet cells were incubated for 30 min at 37°C with the Ca^2+^ indicator Fura-2 AM and then checked for Ca^2+^ influx in response to different modulators (glucose, glibenclamide and diazoxide). Fura-2 AM fluorescence, which ratiometrically indicates cytosolic Ca^2+^ levels, was checked using a Leica DMi8 wide-field fluorescence microscope operated with the LAS X Software (Leica Microsystems), which allows live cell imaging at 37°C in a controlled atmosphere (5% CO_2_, 95% air). Cells were imaged under basal conditions in HBBS buffer with low glucose (5.6 mM) for 5 min, followed by addition of i) high glucose (final 20 mM); ii) 10 µM glibenclamide (K_ATP_ channel blocker); iii) 250 µM diazoxide (K_ATP_ channel opener) + high glucose (final 20 mM). A filter wheel installed in the light source permitted Fura2 fluorescence excitation at 340 nm (Ca^2+^-bound) or 387 nm (Ca^2+^-free). The emission at 510 nm was filtered with a FURA2 filter cube and detected by a DFC365FX-760554215 camera in a binning format of 2×2 (696×520 pixels). Mouse islets were imaged using a HC PL FLUOTAR L 20x/0.40 DRY objective over a time course of approximately 15 min (1 frame each 1.026 s) with exposure times and gain of 600 ms/2 (340 nm) and 50 ms/1 (387 nm). Rat islets were imaged with the same objective for 20 minutes (1 frame each 1.006 s) with exposure times and gain of 600 ms/2 (340 nm) and 30 ms/1 (387 nm). Images were further analyzed using ImageJ FIJI Software.

The steps of this protocol were performed as follows.

1. After cells properly adhere overnight, incubate cells for 30 min with 4 µM of Fura-2 AM at 37°C in complete RPMI medium.
2. With a fluorescence microscope at 37°C, in controlled atmosphere (5% CO_2_, 95% air), register basal fluorescence ratios for at least 5 minutes under low glucose (5.6 mM) conditions.
3. After 5 minutes (basal fluorescence), add the stimulus (physiological or pharmacological) and continue registering the fluorescence for 15-20 minutes. **Note:** The duration may vary depending on the mechanism of action of the stimulus used.

Ca^2+^ influx measurements (Fig. 3-4) demonstrate the ability of dispersed and adhered cells, from both mouse and rat, to respond to physiological and pharmacological stimuli. As seen in Fig. 3-4, cells responded to glibenclamide by rapidly increasing cellular Ca^2+^ influx. Similarly, glucose was also able to increase the Ca^2+^ signal, but in a more gradual manner. The gradual increase in Ca^2+^ in response to glucose, in contrast to the instantaneous effect of glibenclamide, has different possible explanations. While glibenclamide has a direct effect on K_ATP_ channels, glucose needs to be intracellularly metabolized, increasing ATP and then leading to K_ATP_ channel closure. Another possibility may be related to particularities among different beta cell populations, with some groups having a faster response compared to groups with a slower response; since the sample signal comes from the sum of these populations, the increment is gradual. Finally, in the presence of diazoxide, glucose was not able to lead to an increase in Ca^2+^ influx. Our results therefore show that cell dispersion and adhesion is an interesting tool for populational analysis of these cells, isolating the paracrine component from cell-to-cell communication. Furthermore, our results indicate that the expected physiological and pharmacological signaling pathways are preserved in the dispersed islet protocol developed.

**Figure 3.**
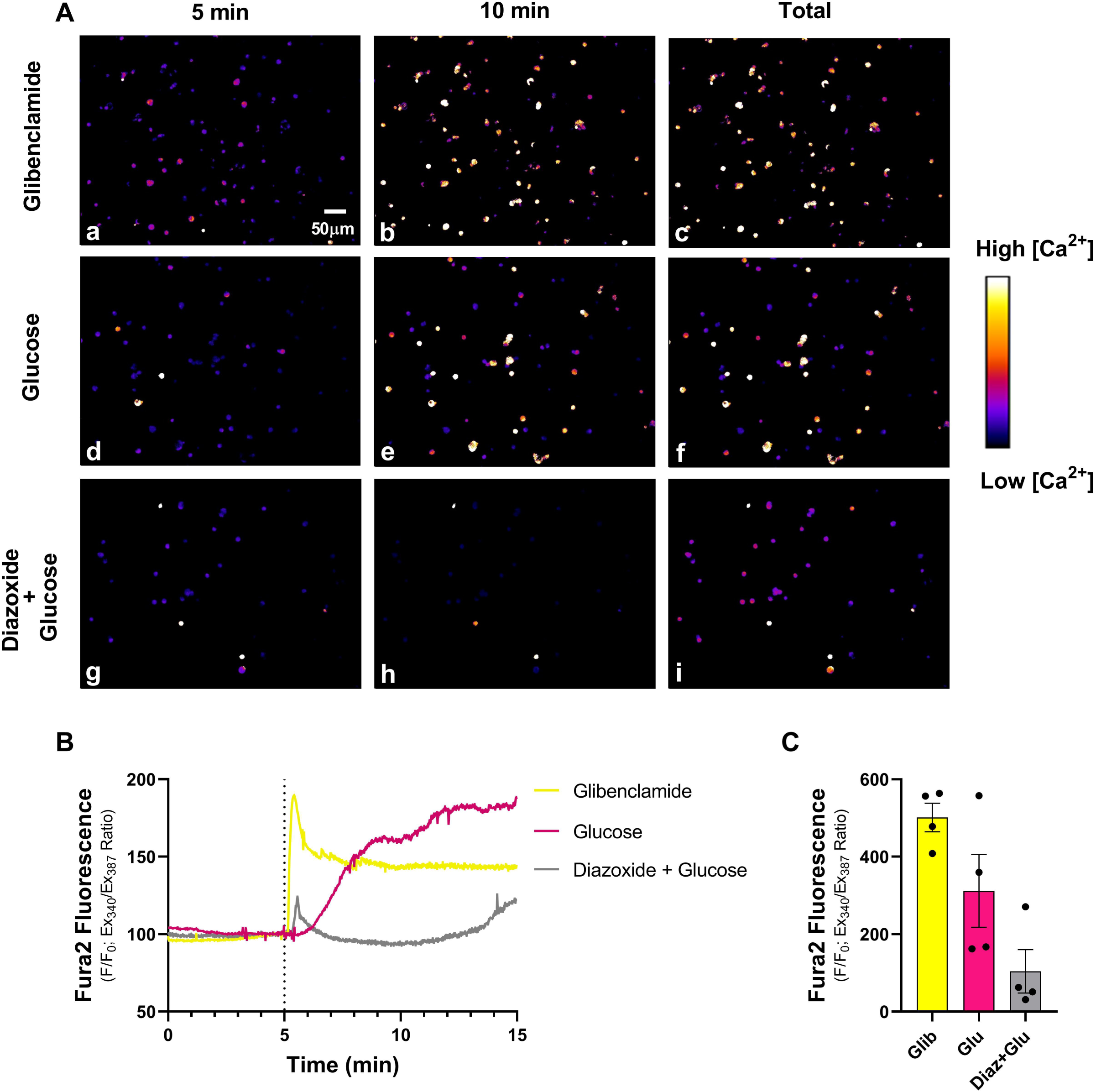
Calcium influx assay in dispersed and adhered mouse islets. Live imaging of dispersed and adhered islet cells under basal conditions with low glucose (5.6 mM), followed by addition of high glucose (final 20 mM); 10 µM glibenclamide; or 250 µM diazoxide + high glucose (20 mM). Representative Fura2-AM fluorescence ratios (340/ 387 nm) in 5 min, 10 min and full trace **(panel A)** under glibenclamide **(a,b,c)**; glucose **(d,e,f)** or diazoxide + high glucose **(g,h,i)**. Fluorescence was measured at baseline for 5 min, followed by different additions as indicated, up to 15 min total. **(B)** Representative Ca^2+^ influx curves. **(C)** Fura2-AM fluorescence ratio (340/ 387 nm) quantifications. Glucose **(pink)**; glibenclamide **(yellow)** and diazoxide + glucose **(grey)**. Results are mean ± SEM of 4 animals.

**Figure 4.**
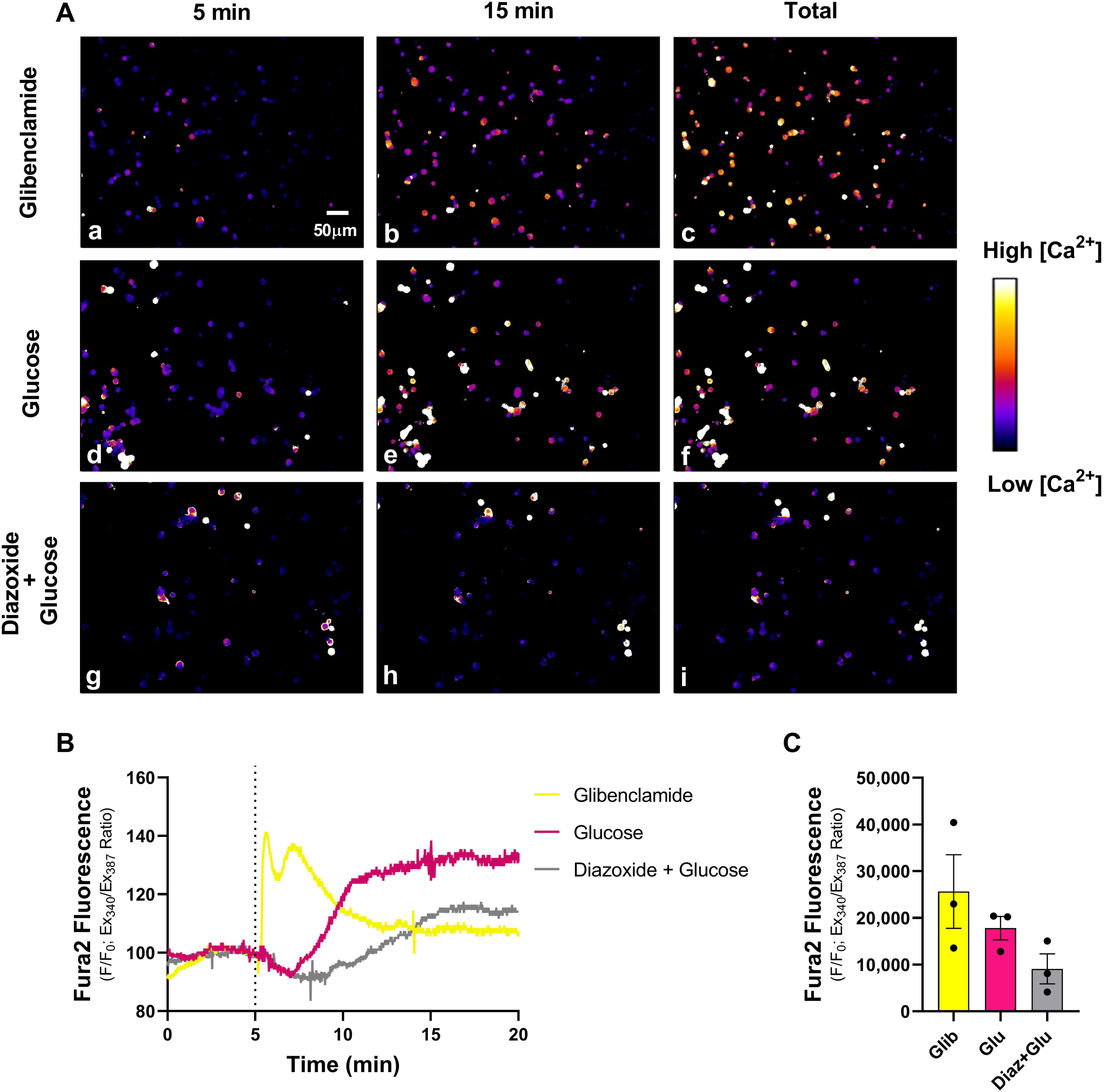
Calcium influx assay in dispersed and adhered rat islets. Live imaging of dispersed and adhered islet cells under basal conditions with low glucose (5.6 mM), followed by addition of high glucose (final 20 mM); 10 µM glibenclamide; or 250 µM diazoxide + high glucose (20 mM). Representative Fura2-AM fluorescence ratios (340/ 387 nm) in 5 min, 15 min and full trace **(panel A)** under glibenclamide **(a,b,c)**; glucose **(d,e,f)** or diazoxide + high glucose **(g,h,i)**. Fluorescence was measured at baseline for 5 min, followed by different additions as indicated, up to 20 min total. **(B)** Representative Ca^2+^ influx curves. **(C)** Fura2-AM fluorescence ratio (340/ 387 nm) quantifications. Glucose **(pink)**; glibenclamide **(yellow)** and diazoxide + glucose **(grey)**. Results are mean ± SEM of 3 animals.

#### 3.4.2. Glucose-stimulated insulin secretion (GSIS) in dispersed islets

The dispersed cells were also checked for GSIS, aiming to evaluate the functionality of the dispersed and adhered islets, as follows:

1. After cells properly adhere overnight, replace culture medium supernatant by 150 µL Krebs Henseleit (KH) buffer (table 2) containing 2.8 mM (for mice) or 5.6 mM glucose (for rats).
2. Preincubate at 37°C for 30 min.
3. Discard the supernatant and replace it with 150 µL KH buffer with low (5.6 mM) or high (20 mM) glucose concentrations.
4. Incubate at 37°C for 60 min.
5. Collect the supernatant in fresh conic tubes and store at -20°C.
6. Remove any remaining buffer and lyse cells with 30 µL of RIPA buffer. **Note**: The samples, insulin-containing supernatants, and plates with RIPA-lysed cells can be stored at -20°C for up to one month.
7. Quantify total protein content using the BCA Pierce protocol.
8. Measure insulin by enzyme-linked immunosorbent assay (ELISA) using a commercial kit (EZRMI-13K, Merck Millipore Corporation, Billerica, USA). Express insulin in ng.ml^-1^. **Note:** Several commercial kits are currently available to measure insulin. Before purchasing the kit, make sure that the range of the standard curve is appropriate for the range of your samples and that you have the volume required for the assay. **Note:** In addition to ELISA, insulin can also be measured by radioimmunoassay, Fluorescence Resonance Energy Transfer (FRET), or High-Performance Liquid Chromatography (HPLC), depending on the availability of these methods for your group.

Evaluating insulin secretion in response to glucose in dispersed and adhered islets, we confirmed that these cells are functional. An increment from 5.6 mM to 20 mM in glucose concentrations was able to increase insulin secretion by 2.5 and 2 times in mouse and rat samples, respectively (Fig. 5), confirming that our protocol provides fully functional islets. Next, we evaluated if the protocol was adequate for metabolic flux analysis.

**Figure 5.**
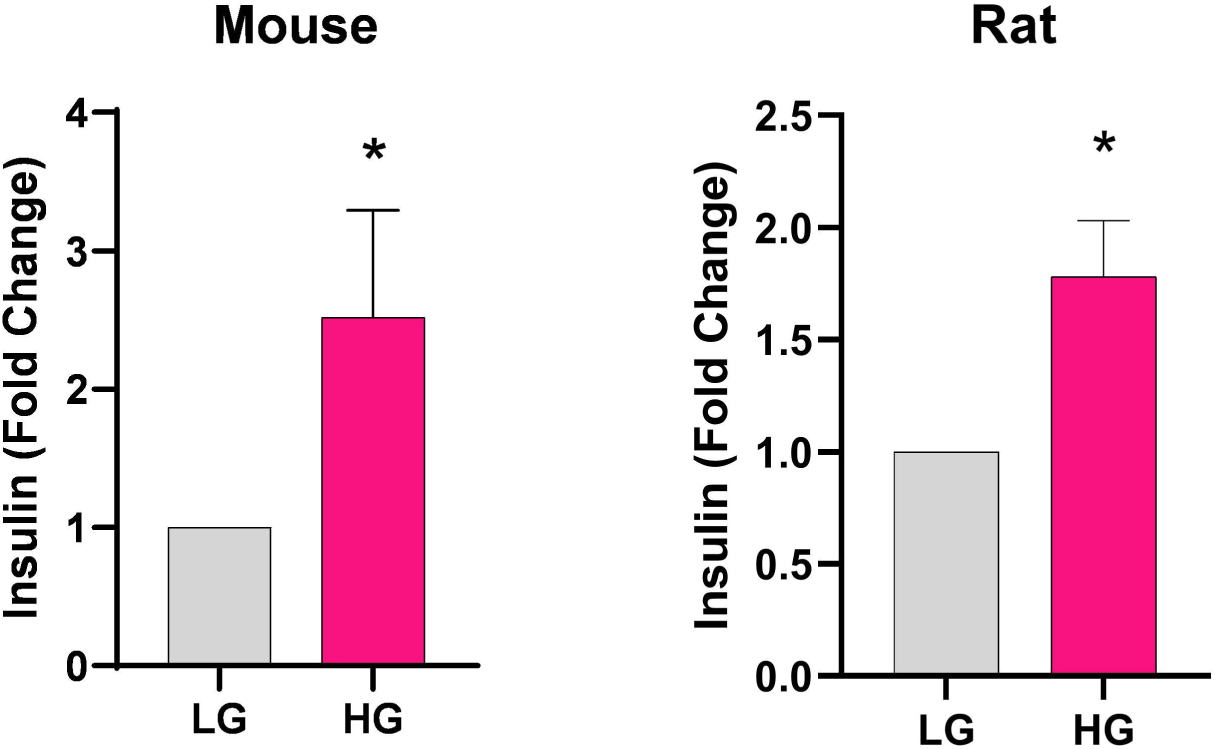
Glucose-stimulated insulin secretion in dispersed and adhered mouse and rat islets. Insulin secretion (Fold change) in response to glucose in dispersed and adhered islets. The increment from 5.6 mM to 20 mM glucose increased insulin secretion from mouse **(A)** and rat cells **(B)**. Results are mean ± SEM of 5 mice and 4 rats. *p<0.05 using paired Student’s t test.

### 3.5. Oxygen consumption in dispersed islets

After overnight incubation on Agilent Seahorse XF24 cell culture microplates previously coated with poly-L-lysine, oxygen consumption was measured using a Seahorse Extracellular Flux Analyzer (Agilent Technologies, Santa Clara, CA, EUA).

**Note:** Remember to include cell-free wells with the same medium in the microplates for background correction.

1. One day prior to the assay: hydrate the Seahorse cartridge with appropriate calibrant solution overnight at 37°C in a humidified incubator without CO_2_.
2. Replace RPMI complete medium by 500 µL RPMI containing 1% penicillin/streptomycin (P/S) and 5 mM HEPES (without FBS nor bicarbonate). **Note:** It is also possible to use the commercially available Seahorse medium (without FBS nor bicarbonate).
3. Incubate cells for 1 h at 37°C in a humidified incubator without CO_2_.
4. Prepare working solutions and pipette solutions at each port.
5. Set the equipment: select different groups and different injections according to the protocol. **Note:** Remember to select cell-free wells for background correction. **Note:** To avoid cell detachment, we adjusted the regular timepoints to 1 min mixing + 2 min waiting + 3 min of measurements. **Note:** We tested two protocols:
  i. Basal respiration at 10 mM glucose, followed by the standard Mito stress test [28].
  ii. Basal respiration at 5.6 mM glucose, addition of high glucose at port A (final concentration: 20 mM), followed by the Mito stress test.
  **Note:** Mito stress test injections: a) oligomycin (final concentration: 2 µM); b) CCCP (final concentration: 6 µM); c) rotenone and antimycin (R/AA, final concentration: 1 µM each).
6. Place the cartridge in the Seahorse equipment for appropriate system calibration.
7. After 1 h incubation of cells and once calibration is ready, remove the microplate with the calibrant and place the microplate with cells in the equipment to measure OCR.
8. At the end of each experiment, remove the medium carefully and lyse cells with 30 µL RIPA buffer per well.
9. Quantify total protein using the BCA Pierce protocol. Normalize OCR values obtained per amount of total protein in each well. **Note:** Within each experiment, selecting islets of the same size decreases the variability between replicates. However, there are variations between animals, because different animals have islets of different sizes. **Note:** We used the XF24 analyzer, as this is the available equipment in our facility, but our method, in principle, could be adapted for all Seahorse analyzers, including 96 and 6 well apparatuses.

We find that both sample origins (mouse and rat) present the expected response to mitochondrial modulators (oligomycin, CCCP and antimycin + rotenone) (Fig. 6A,B and 7A,B), as we observe a decrease in OCR after oligomycin addition (which inhibits ATP synthase), an increase after CCCP (which reduces the mitochondrial inner membrane potential, maximizing electron transport) and an expressive decrease after the injection of antimycin/rotenone (which inhibit electron transport). In response to glucose, we observe an expected and significant increase in OCR, which is more evident in mouse islets (Fig. 6C and D) compared to rats (Fig. 7C and D). We suggest that this difference may be due to the number of islets used (20 for rats and 40 for mice). However, and most importantly, the response is still clearly observed in both cases.

**Figure 6.**
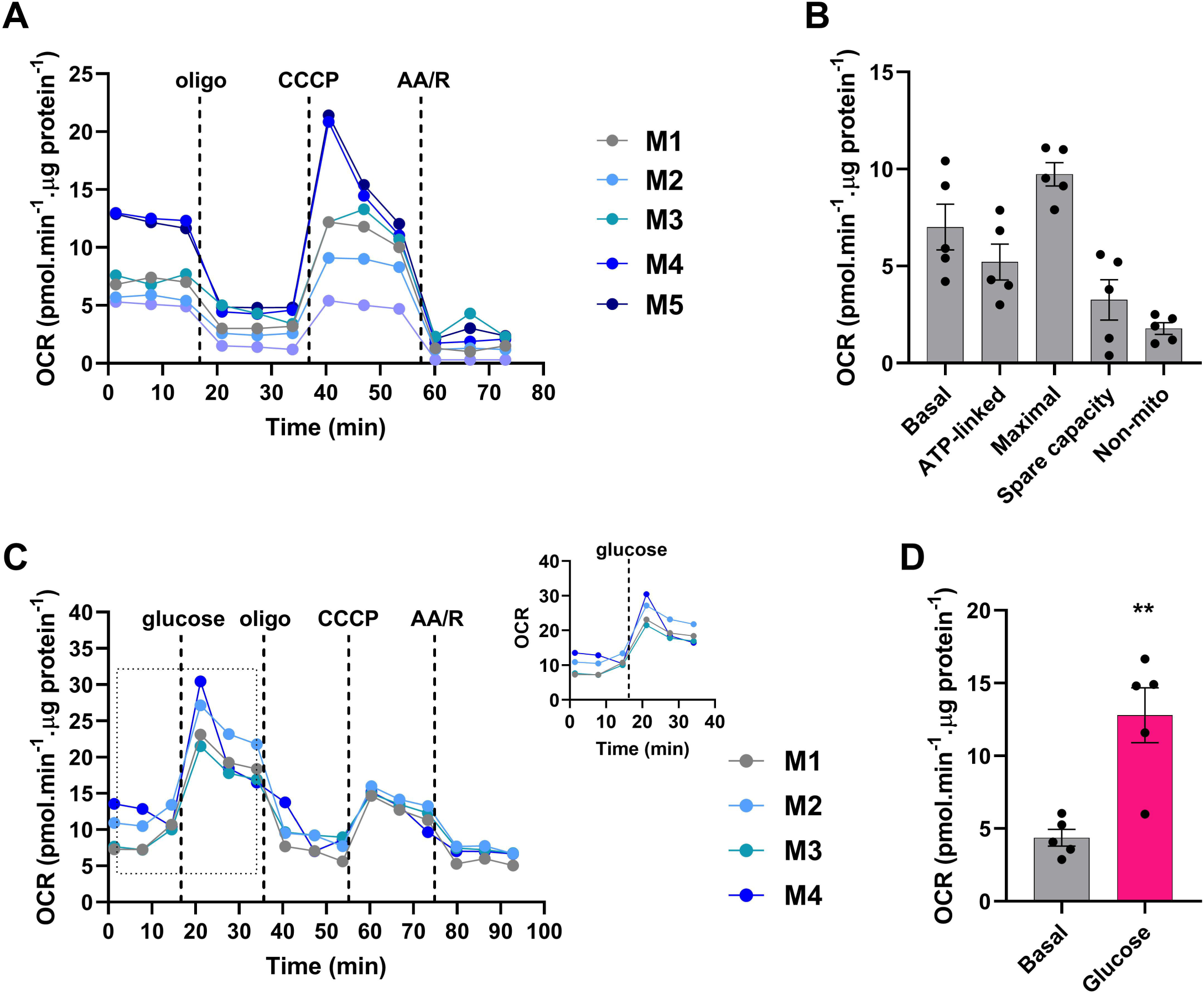
Oxygen Consumption Rates (OCR) in dispersed and adhered mouse islets. **(A)** OCR curves of 40 dispersed mouse islets in response to mitochondrial modulators oligomycin (2 µM), CCCP (6 µM) and Antimycin+Rotenone (1 µM each). **(B)** Quantification of mean basal, ATP-linked, maximal, and non-mitochondrial respiration. **(C)** OCR curves in response to glucose and mitochondrial modulators. Insets represent the magnified area indicated by the dotted lines. **(D)** Quantification of the mean OCR value. Results are mean ± SEM of 5 **(A-B)** or 4 **(C-D)** mice. M1-M5 represent the values of individual animals used. **p<0.01 using paired Student’s t test.

**Figure 7.**
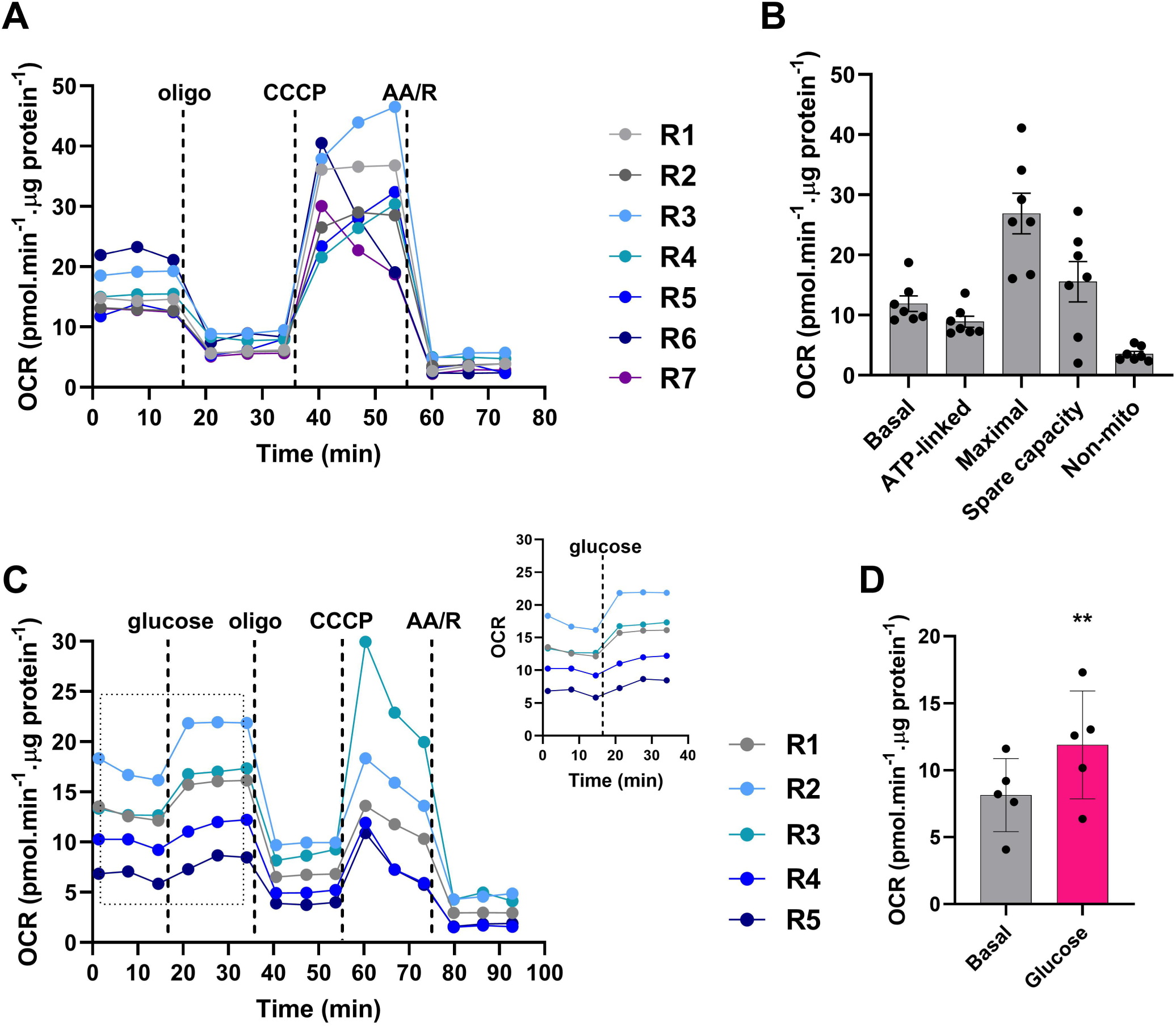
Oxygen Consumption Rates (OCR) in dispersed and adhered rat islets. **(A)** OCR curves of 20 dispersed rat islets in response to mitochondrial modulators oligomycin (2 µM), CCCP (6 µM) and Antimycin+Rotenone (1 µM each). **(B)** Quantification of mean basal, ATP-linked, maximal, and non-mitochondrial respiration. **(C)** OCR curves in response to glucose and mitochondrial modulators. Insets represent the magnified area indicated by the dotted lines. **(D)** Quantification of the mean OCR value. Results are mean ± SEM of 7 **(A-B)** or 5 **(C-D)** rats. R1-R7 represent the values of individual animals used. **p<0.01 using paired Student’s t test.

Interestingly, the prior addition of glucose decreases the maximum OCRs notably in both mouse (Fig. 6C) and rat samples (Fig. 7C). This may be expected, since these cells respond to glucose by promoting overt increases in Ca^2+^ influx, which at high levels at the mitochondrial matrix may limit electron transport [29, 30]. As a result, there is a decrease in the response to the uncoupler CCCP. This indicates that experimental designs should be developed specifically for different desired readouts: experiments in which maximized OCRs induced by uncouplers are important parameters should be conducted without a high glucose stimulus prior to the addition of CCCP.

## 4. Dis(advantages) of different methods using the Seahorse Extracellular Flux System

As stated above, mitochondrial oxygen consumption can be measured mainly with high resolution using two commercial setups: Oroboros Respirometers and Seahorse Extracellular Flux Analyzers. Although Oroboros Oxygraphy is a robust method, it requires a high number of islets per condition, which means more than one animal per condition. Without the specific chamber to contain the islets, it also compromises islet viability throughout the measurements due to mechanical damage. Therefore, we consider the Seahorse Extracellular Flux System a more suitable method for islets. In Table 1, we compare the different Seahorse methods available with ours.

The specific islet microplate keeps the islets trapped underneath a grid, preventing, although not completely, them from floating away during the measurements. On the other hand, it is more expensive than the standard microplate, requires a specific tool to position the grid on top of the wells and is only compatible with the XF24 system, in addition to being laborious and time consuming, with considerable variations between experiments. The spheroid microplate keeps the islets adhered when a proper precoating is used, although it does not guarantee that islets do not detach during measurements, leading to variations between replicates. It is more expensive than the standard microplate, is only compatible with the XF96 system, and needs a specific thermal tray. We propose a new and practical method using dispersed islets and the standard microplate (Fig. 8), which is compatible with both XF24 and XF96 systems and does not need additional consumables. Our method was developed for rodent islets, but in principle it can also be used for human islets. In addition, this method also allows microscopy assays. This way, particularities between different endocrine cell types can be better explored, since the resolution of the microscope makes it possible to assess differences at the single cell level.

**Figure 8.**
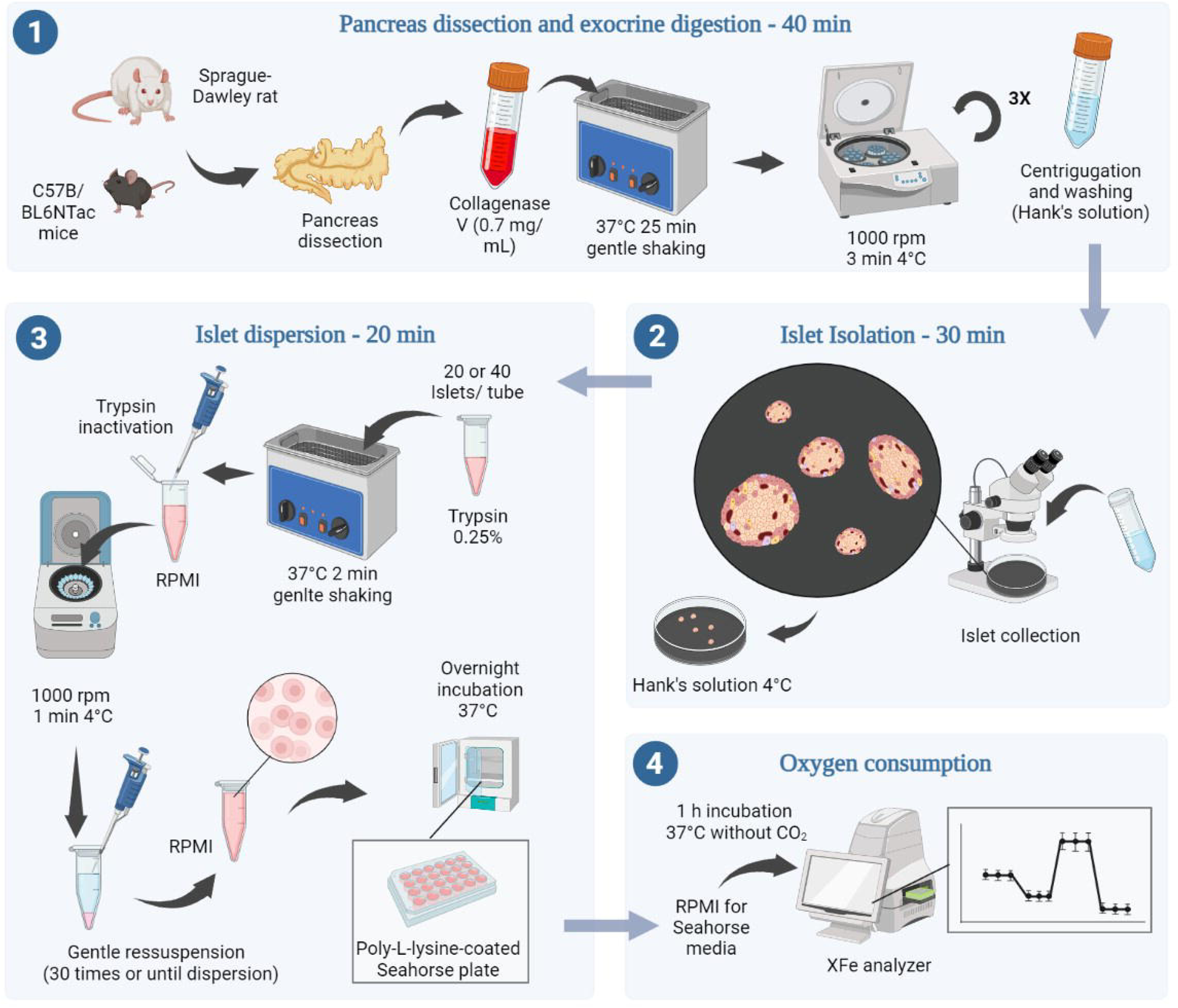
A practical method to evaluate *ex vivo* dispersed islet respiration. Schematic workflow of all steps necessary to isolate, disperse, and culture mouse and rat islets, followed by oxygen consumption rate assays. Created with BioRender.com.

## 5. Buffers and culture media composition

### 5.1. Hanks’ Balanced Salt Solution (HBSS) - used for islet isolation

1. Add 800 mL of milliQ water to a suitable container;
2. Add NaCl, KCl, MgSO_4_, Na_2_HPO_4_ and KH_2_PO_4_, according to table 1;
3. Adjust pH to 6.3;
4. Add CaCl_2_, NaHCO_3_, D-glucose, bovine serum albumin, penicillin and streptomycin (P/S);
5. Adjust pH to 7.4 and final volume to 1 L;
6. Filter the solution using a 0.2 µm filter to a sterile flask;
7. Store it at 4°C until use.

### 5.2. Krebs Henseleit (KH) buffer - used for islet isolation

1. Mix equal volumes of each solution (I, II + III) with same volume of milliQ water;
2. Adjust pH to 7.4, using bubbling carbogen. OBS.: do not use acid or base to adjust pH.
3. Add 0.2 % BSA (2 g for a final 1 L).

### 5.2. RPMI 1640 culture medium

1. Supplement with bicarbonate (2 g/L), 10% FBS (100 mL for a final 1 L) and 1% P/S.
2. Adjust pH to 7.4 and filter using a 0.2 µm filter.

**Note:** For cell culture, we used media with 11.1 mM glucose and phenol red (#31800-022). For cell imaging, we used a media with 5.6 mM glucose and without phenol red (#103576-100).

### 5.4. Hanks’ Balanced Salt Solution (HBSS) - used for Ca_2+_ imaging (#14175-095, Sigma)

1. Supplement with 2 mM CaCl_2_ and 0.4 mM MgSO_4_. To a final 30 mL HBSS, add 60 µL CaCl_2_ 1 M and 12 µL MgSO_4_ 1 M.
2. Adjust pH to 7.4.

## 6. Troubleshooting

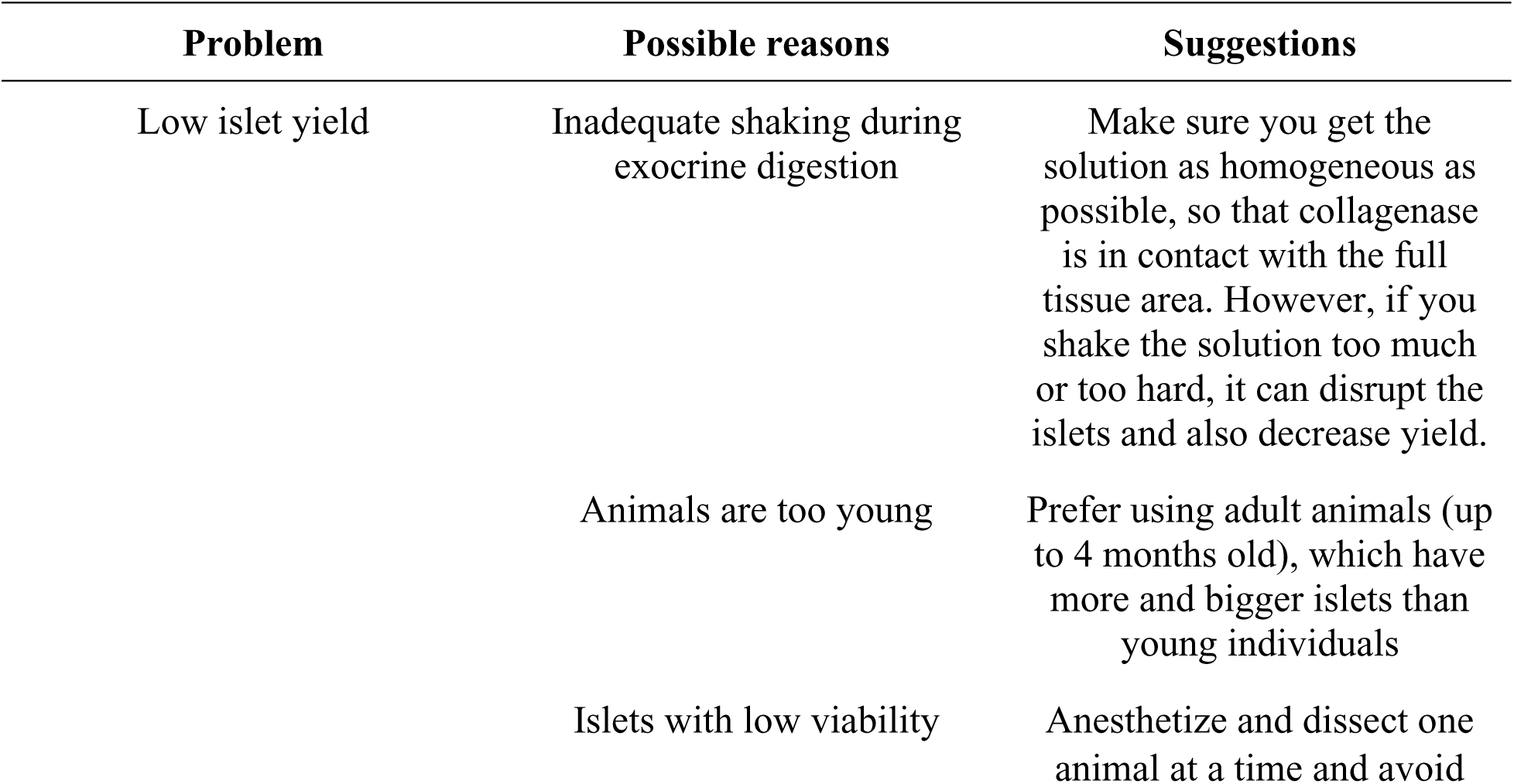
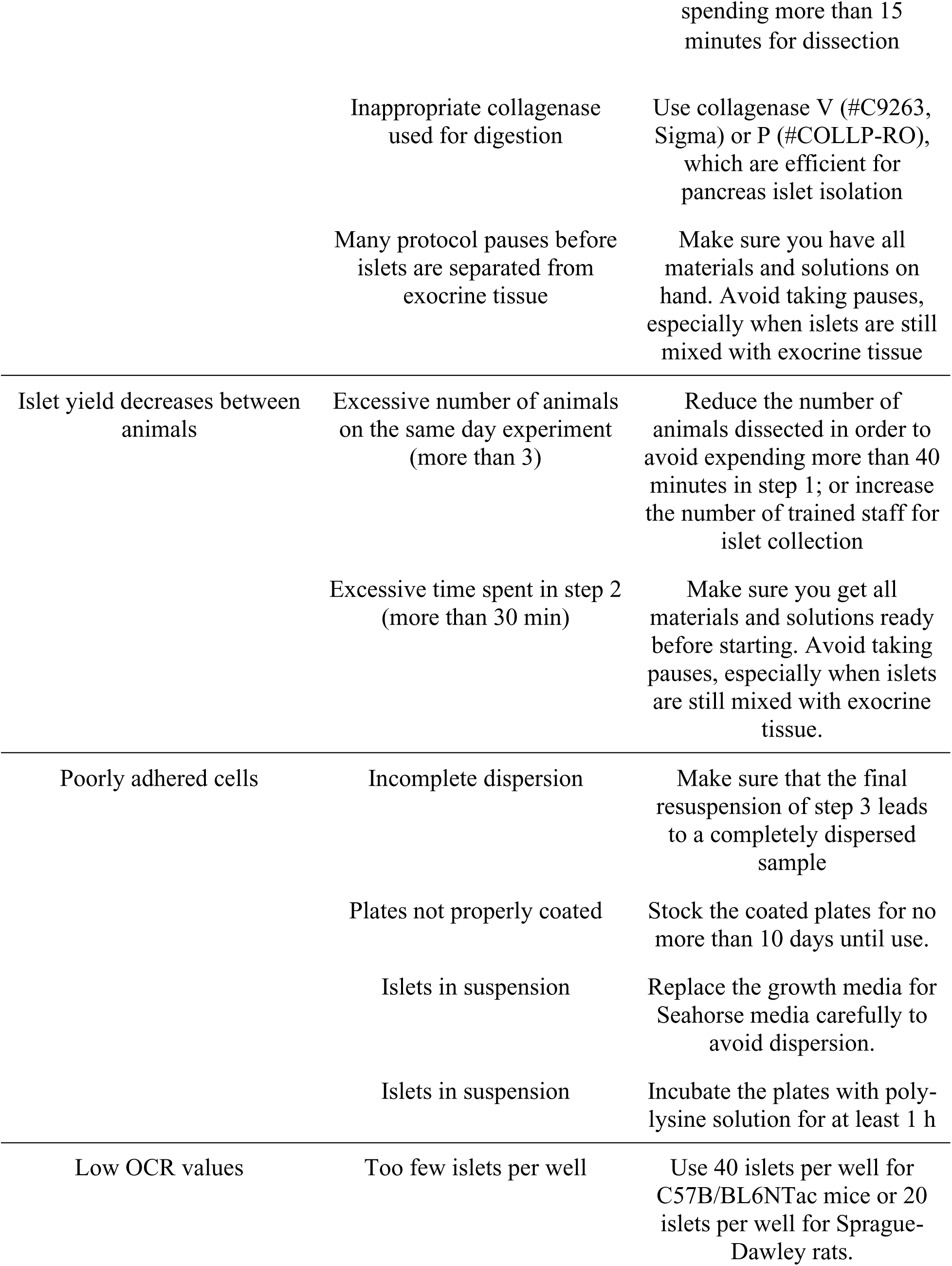
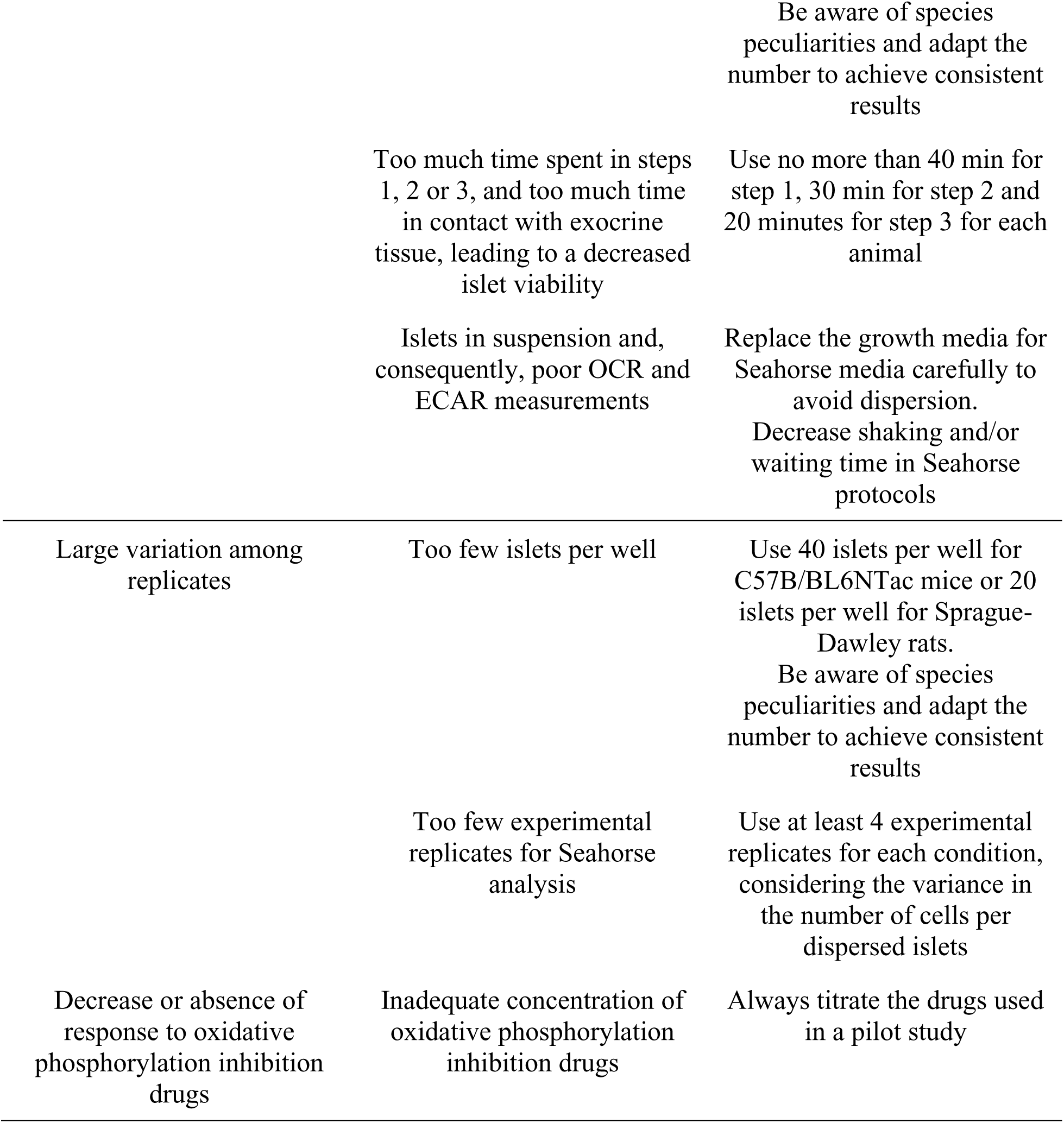

## Acknowledgements

The authors thank Camille C. Caldeira da Silva for excellent technical assistance. We also acknowledge Silvânia M. P. Neves and her animal facility crew for exceptional expert animal care.

## Data availability

All data generated or analyzed during this study are included in this published article.

## Authors’ relationships and activities

The authors declare that there are no relationships or activities that might bias, or be perceived to bias, their work.

## Contribution statement

EAVB and DSR are responsible for conception and design of research; EAVB, DSR and ACM performed experiments and analyzed the data; ABC provided instrumentation and participated in formal analysis; EAVB and DSR interpreted results of experiments and prepared figures; EAVB, DSR and AJK drafted the manuscript; all authors revised and approved the final version of the manuscript. EAVB is the guarantor of this work and, as such, had full access to all the data in the study and takes responsibility for the integrity of the data and the accuracy of the data analysis.

## References

[1] Huang HH, Harrington S, Stehno-Bittel L (2018) The Flaws and Future of Islet Volume Measurements. Cell Transplant 27(7): 1017–1026. 10.1177/0963689718779898

[2] Kilimnik G, Jo J, Periwal V, Zielinski MC, Hara M (2012) Quantification of islet size and architecture. Islets 4(2): 167–172. 10.4161/isl.19256

[3] Kim A, Miller K, Jo J, Kilimnik G, Wojcik P, Hara M (2009) Islet architecture: A comparative study. Islets 1(2): 129–136. 10.4161/isl.1.2.9480

[4] Steiner DJ, Kim A, Miller K, Hara M (2010) Pancreatic islet plasticity: interspecies comparison of islet architecture and composition. Islets 2(3): 135–145. 10.4161/isl.2.3.11815

[5] Rorsman P, Ashcroft FM (2018) Pancreatic β-Cell Electrical Activity and Insulin Secretion: Of Mice and Men. Physiol Rev 98(1): 117–214. 10.1152/physrev.00008.2017

[6] Wiederkehr A, Wollheim CB (2012) Mitochondrial signals drive insulin secretion in the pancreatic β-cell. Mol Cell Endocrinol 353(1-2): 128–137. 10.1016/j.mce.2011.07.016

[7] Maechler P (2013) Mitochondrial function and insulin secretion. Mol Cell Endocrinol 379(1-2): 12–18. 10.1016/j.mce.2013.06.019

[8] Anello M, Lupi R, Spampinato D, et al. (2005) Functional and morphological alterations of mitochondria in pancreatic beta cells from type 2 diabetic patients. Diabetologia 48(2): 282–289. 10.1007/s00125-004-1627-9

[9] Chen J, Stimpson SE, Fernandez-Bueno GA, Mathews CE (2018) Mitochondrial Reactive Oxygen Species and Type 1 Diabetes. Antioxid Redox Signal 29(14): 1361–1372. 10.1089/ars.2017.7346

[10] Yamauchi Y, Nakamura A, Yokota T, et al. (2022) Luseogliflozin preserves the pancreatic beta-cell mass and function in db/db mice by improving mitochondrial function. Sci Rep 12(1): 9740. 10.1038/s41598-022-13888-6

[11] Crowder JJ, Zeng Z, Novak AN, Alves NJ, Linnemann AK (2022) Stabilization protects islet integrity during respirometry in the Oroboros Oxygraph-2K analyzer. Islets 14(1): 128–138. 10.1080/19382014.2022.2054251

[12] Regeenes R, Wang Y, Piro A, et al. (2023) Islet-on-a-chip device reveals first phase glucose-stimulated respiration is substrate limited by glycolysis independent on Ca2+ activity. In, Biosensors and Bioelectronics: X

[13] Taddeo EP, Stiles L, Sereda S, et al. (2018) Individual islet respirometry reveals functional diversity within the islet population of mice and human donors. Mol Metab 16: 150–159. 10.1016/j.molmet.2018.07.003

[14] Fukunaka A, Shimura M, Ichinose T, et al. (2023) Zinc and iron dynamics in human islet amyloid polypeptide-induced diabetes mouse model. Sci Rep 13(1): 3484. 10.1038/s41598-023-30498-y

[15] Kim YK, Walters JA, Moss ND, et al. (2022) Zinc transporter 8 haploinsufficiency protects against beta cell dysfunction in type 1 diabetes by increasing mitochondrial respiration. Mol Metab 66: 101632. 10.1016/j.molmet.2022.101632

[16] Wikstrom JD, Sereda SB, Stiles L, et al. (2012) A novel high-throughput assay for islet respiration reveals uncoupling of rodent and human islets. PLoS One 7(5): e33023. 10.1371/journal.pone.0033023

[17] Coltman NJ, Rochford G, Hodges NJ, Ali-Boucetta H, Barlow JP (2022) Exploring Mitochondrial Energy Metabolism of Single 3D Microtissue Spheroids Using Extracellular Flux Analysis. J Vis Exp(180). 10.3791/63346

[18] Lacy PE, Kostianovsky M (1967) Method for the isolation of intact islets of Langerhans from the rat pancreas. Diabetes 16(1): 35–39. 10.2337/diab.16.1.35

[19] Corbin KL, West HL, Brodsky S, Whitticar NB, Koch WJ, Nunemaker CS (2021) A Practical Guide to Rodent Islet Isolation and Assessment Revisited. Biol Proced Online 23(1): 7. 10.1186/s12575-021-00143-x

[20] Salvalaggio PR, Deng S, Ariyan CE, et al. (2002) Islet filtration: a simple and rapid new purification procedure that avoids ficoll and improves islet mass and function. Transplantation 74(6): 877–879. 10.1097/00007890-200209270-00023

[21] Li DS, Yuan YH, Tu HJ, Liang QL, Dai LJ (2009) A protocol for islet isolation from mouse pancreas. Nat Protoc 4(11): 1649–1652. 10.1038/nprot.2009.150

[22] Komatsu H, Cook C, Wang CH, et al. (2017) Oxygen environment and islet size are the primary limiting factors of isolated pancreatic islet survival. PLoS One 12(8): e0183780. 10.1371/journal.pone.0183780

[23] Komatsu H, Kandeel F, Mullen Y (2018) Impact of Oxygen on Pancreatic Islet Survival. Pancreas 47(5): 533–543. 10.1097/MPA.0000000000001050

[24] Weir GC, Halban PA, Meda P, Wollheim CB, Orci L, Renold AE (1984) Dispersed adult rat pancreatic islet cells in culture: A, B, and D cell function. Metabolism 33(5): 447–453. 10.1016/0026-0495(84)90146-x

[25] Scarl RT, Koch WJ, Corbin KL, Nunemaker CS (2021) Isolation and Assessment of Pancreatic Islets Versus Dispersed Beta Cells: A Straightforward Approach to Examine Cell-Cell Communication. Methods Mol Biol 2346: 151–164. 10.1007/7651_2020_338

[26] Hartig SM, Cox AR (2020) Paracrine signaling in islet function and survival. J Mol Med (Berl) 98(4): 451–467. 10.1007/s00109-020-01887-x

[27] Félix-Martínez GJ, Godínez-Fernández JR (2023) A primer on modelling pancreatic islets: from models of coupled β-cells to multicellular islet models. Islets 15(1): 2231609. 10.1080/19382014.2023.2231609

[28] Gu X, Ma Y, Liu Y, Wan Q (2021) Measurement of mitochondrial respiration in adherent cells by Seahorse XF96 Cell Mito Stress Test. STAR Protoc 2(1): 100245. 10.1016/j.xpro.2020.100245

[29] Halestrap AP (2006) Calcium, mitochondria and reperfusion injury: a pore way to die. Biochem Soc Trans 34(Pt 2): 232–237. 10.1042/BST20060232

[30] Vilas-Boas EA, Cabral-Costa JV, Ramos VM, Caldeira da Silva CC, Kowaltowski AJ (2023) Goldilocks calcium concentrations and the regulation of oxidative phosphorylation: Too much, too little, or just right. J Biol Chem 299(3): 102904. 10.1016/j.jbc.2023.102904

